# 3D printed magneto-active microfiber scaffolds for remote stimulation of 3D *in vitro* skeletal muscle models

**DOI:** 10.1101/2023.01.19.524679

**Authors:** Gerardo Cedillo-Servin, Ouafa Dahri, João Meneses, Joost van Duijn, Fanny Sage, Joana Silva, André Pereira, Fernão D. Magalhães, Jos Malda, Niels Geijsen, Artur M. Pinto, Miguel Castilho

## Abstract

Tunable culture platforms that guide cellular organization and mechanically stimulate skeletal muscle development are still unavailable due to limitations in biocompatibility and actuation triggered without contact. This study reports the rational design and fabrication of magneto-active microfiber meshes with controlled hexagonal microstructures via melt electrowriting (MEW) of a thermoplastic/graphene/iron oxide composite. *In situ* deposition of iron oxide nanoparticles on oxidized graphene yielded homogeneously dispersed magnetic particles with sizes above 0.5 μm and low aspect ratio, preventing cellular internalization and toxicity. With these fillers, homogeneous magnetic composites with very high magnetic filler content (up to 10 wt.%) were obtained and successfully processed in a solvent-free manner for the first time. MEW of magnetic composites enabled the skeletal muscle-inspired design of hexagonal scaffolds with tunable fiber diameter, reconfigurable modularity, and zonal distribution of magneto-active and nonactive material. Importantly, the hexagonal microstructures displayed elastic deformability under tension, mitigating the mechanical limitations due to high filler content. External magnetic fields below 300 mT were sufficient to trigger out-of-plane reversible deformation leading to effective end-to-end length decrease up to 17%. Moreover, C2C12 myoblast culture on 3D Matrigel/collagen/MEW scaffolds showed that the presence of magnetic particles in the scaffolds did not significantly affect viability after 8 days with respect to scaffolds without magnetic filler. Importantly, *in vitro* culture demonstrated that myoblasts underwent differentiation at similar rates regardless of the presence of magnetic filler. Overall, these innovative microfiber scaffolds were proven as a magnetically deformable platform suitable for dynamic culture of skeletal muscle with potential for *in vitro* disease modeling.

## 1. Introduction

Skeletal muscle is the most abundant tissue in the human body (40–50 mass %) and is essential for posture, locomotion, and physiological and metabolic processes ^[1]^. Proper skeletal muscle function is highly dependent on the synergy among three interconnecting structures separated by extracellular matrix (ECM): the epimysium —surrounding the entire muscle bundle–, the perimysium –surrounding multiple muscle fibers within a muscle—, and the endomysium, which surrounds each muscle fiber individually ^[2]^. This unique organization provides mechanical support to the hierarchical muscle structures and transmits contractile forces from the myocytes within the muscle bundle ^[3,4]^.

In the event of genetic defects in sarcolemmal, contractile, or ECM proteins, skeletal muscle function is rapidly lost leading to muscular dystrophies (MD), such as facioscapulohumeral dystrophy (FSHD) and Duchenne MD (DMD). MD are known to involve progressive weakness that severely impacts patients’ lives by causing disability and, ultimately, death due to cardiac or respiratory failure. In particular, DMD is an X chromosome-linked condition affecting about one in 5,000 males yearly, and most patients require assisted ventilation at around 20 years of age ^[5,6]^; meanwhile, about 10% of FSHD patients eventually become wheelchair dependent ^[7]^.

Current available approaches to study MD both *in vitro* and *in vivo* are limited ^[5,8]^. Differentiation and functional maturation of skeletal muscle progenitor cells is commonly achieved using growth factors and small molecules, and mechanical stimulation further contributes to maintaining the structure of adult skeletal muscle ^[8]^. For example, cyclical mechanical stimulation of myoblasts alone has been shown to enhance expression of myogenic differentiation markers ^[9]^. Therefore, the efficacy of *in vitro* models of MD can greatly benefit from 3D matrices that restore myofiber organization by introducing anisotropic mechanical environments and, importantly, support and actively stimulate muscular contractile forces during regeneration ^[10]^. Moreover, engineered skeletal muscle for disease modeling of MD would benefit from deformable matrices that resemble the native extracellular matrix. The microenvironment of skeletal muscle is composed of a 3D network of hierarchically organized fibers which are key in force generation and orientation by providing critical topographical cues and spatial boundary conditions to the cells^[8]^. For instance, synthetic matrices produced by additive manufacturing are able to guide myocyte alignment^[11]^ and have been observed to improve expression of myogenic genes and maturation^[12]^.

Fiber-based scaffolds that mimic skeletal muscle structure and function have offered a promising strategy to guide cell alignment and therefore cell organization. For instance, alginate and gelatin hydrogel-coated millimeter-thick fibers produced by wet spinning have enabled the fabrication of myocyte-laden constructs ^[13]^, while aligned electrospun poly(lactide-*co*-glycolide) (PLGA) fiber scaffolds have been shown to promote the formation of myofiber networks *in vivo* following implantation of a myoblast-seeded scaffold into a murine DMD model ^[14]^. However, existing fiber scaffolds produced by conventional fabrication strategies fail to perform under physiological deformations and, importantly, cannot stimulate muscular contractile forces. This has led to immaturity of transferred cells and limited contractility of newly formed tissue, thus hampering potential clinical use of these strategies. The introduction of fiber processing technologies that allow fabrication of well-organized fiber scaffolds has provided new perspectives for muscle engineering both *in vitro* and *in vivo* ^[10]^.

Melt electrowriting (MEW) is a technology that combines additive manufacturing principles with electrohydrodynamic printing, offering an alternative for highly controlled microfiber deposition. 3D constructs often consisting of polycaprolactone (PCL) —the bioresorbable gold standard for MEW— are fabricated by precise, successive fiber-by-fiber stacking *via* continuous solvent-free deposition of fibers with path geometries that can mitigate the intrinsic mechanical limitations of the printed materials. As shown previously with cardiac cells, PCL MEW scaffolds with rectangular pores embedded in hydrogels can promote cell alignment along the rectangle main axis ^[15]^. Moreover, PCL MEW interconnected sinusoidal and hexagonal meshes have also been designed to unlock even further anisotropic control of cardiac cell orientation and support large reversible deformations due to their controlled fiber geometries ^[10,16]^. Despite their great potential, existing MEW scaffolds alone have not been able to mechanically condition muscle cells in a responsive manner, so that active stimulation accelerates or primes regeneration.

Diverse stimuli-responsive mechanisms have been established in soft robotics to mechanically stimulate tissues, including electro-induced ^[17]^ and magnetothermal actuation ^[18]^. In particular, magnetically active biomaterials represent a contactless platform with no risks associated to high electrical or heat leaks. Iron oxide nanoparticles (ION) are a magnetic filler that has been incorporated into thermoplastics and hydrogels for producing magnetized scaffolds ^[19–22]^. However, iron particles have shown cyto- and genotoxicity even at low concentrations (μg mL^- 1^) and with surface coatings ^[19,23]^. The small size of ION –in the order of tens of nanometers– promotes nonspecific internalization and intracellular accumulation, so to avoid toxicity, ION-based magnetoactive biomaterials are limited to low ION content and small magnetically triggered deformations ^[19,22,24]^. Thus, immobilization on other materials has been proposed to decrease cellular internalization ^[25]^. Graphene-based materials (GBM) are promising particles with larger sizes –hundreds to thousands of nanometers– for reduced internalization, showing good biocompatibility at high concentrations *in vitro* and *in vivo*, as well as biodegradation by neutrophils or the human enzyme myeloperoxidase ^[26,27]^. In particular, reduced graphene oxide (rGO) is a popular GBM with greater hemocompatibility than graphene oxide ^[28,29]^ that has been shown to promote myogenic differentiation with low reduction in cell viability ^[13]^.

Previously in the context of MEW, GBM and ION have only been composited with PCL separately, and primarily with organic solvent-based methods, reaching up to 1% content of rGO ^[28]^ or up to 0.3% of ION ^[30]^. In these composite formation approaches, high particle concentrations tend to have low processability due to particle aggregation and high mechanical stifening. Here we assess the use of rGO not as a mechanical reinforcement for PCL, but as a platform for ION deposition that enables organic solvent-free blending and MEW of magneto-active PCL. We hypothesized that ION deposited on rGO could yield magnetic particles that can be processed by MEW after melt-blending with PCL, and that well-organized magnetic PCL fiber scaffolds could be fabricated with a variety of designs for guiding myofiber organization and supplying a remotely controlled platform for mechanical stimulation of myocyte cultures *in vitro*. To test these hypotheses, we first investigated if Fe^2+^ cations in FeCl_2_ could be oxidized by oxidized graphene nanoplatelets (GNP-ox), leading to the deposition of ION *in situ* on the surface of self-reduced graphene nanoplatelets (rGNP) and yielding microscale magnetized powders that bypass the biological toxicity of nanometer-sized magnetic nanoparticles. Then the homogeneous dispersion of magnetic reduced GNP (rGNP@) in a medical-grade thermoplastic PCL matrix was investigated by melt-blending. With this approach, the fabrication of magnetized fiber meshes was investigated by MEW. Magnetic moment changes after processing, printing accuracy, out-of-plane actuation, and mechanical properties of printed scaffolds were rigorously assessed. Moreover, it was evaluated whether MEW allowed for the controlled deposition of zonally distributed materials with customized geometries, thus facilitating the skeletal muscle-inspired design of actuating scaffolds with promise for recapitulating the organization of muscle tissue cells and ECM. Finally, to assess the biological potential of these magnetoactive scaffolds, myoblasts were seeded and cultured *in vitro* on scaffolds composed of magnetic MEW meshes embedded in collagen/Matrigel hydrogels and assessed for viability and differentiation performance.

## 2. Materials and Methods

### Materials

Granular medical-grade poly-(ε-caprolactone) (PCL) (Purasorb PC 12) was purchased from Purac Biomaterials (the Netherlands). Graphene nanoplatelets (GNP) grade C750 were acquired from XG Sciences (United States), with average thickness under 2 nm, surface area of 750 m^2^g^-1^, and platelet length under 2 μm, according to the manufacturer. Iron (II) chloride tetrahydrate (FeCl_2_·4H_2_O, 98.0%, Honeywell Fluka) and ammonium hydroxide (NH_4_OH, 28.0-30.0% NH_3_, Alfa Aesar) were purchased from ThermoFisher Scientific (United States). Filters with 5-8/17-30 μm of pore size were acquired from Filtres Fioroni (France). All materials were used as received unless otherwise stated.

### GNP oxidization and magnetization

GNP were oxidized by the modified Hummers method (MHM), as described elsewhere ^[31]^. Briefly, 320 mL of H_2_SO_4_ and 80 mL of H_3_PO_4_ were added to 8 g of GNP at room temperature and the solution cooled in an ice bath, followed by the gradual addition of 48 g of KMnO_4_. Then 1200 mL of distilled water were gradually added, followed by addition of H_2_O_2_ until oxygen release stopped. Oxidized GNP (GNP-ox) were washed 5 times with distilled water by centrifugation at 4000 rpm for 15 min. Deposition of iron oxide nanoparticles (ION) on GNP-ox was performed as described previously with some modifications ^[32]^. Briefly, water dispersions of FeCl_2_·4H_2_O (40 mg mL^-1^) and GNP-ox (2 mg mL^-1^) dispersions in water were mixed and sonicated for 10 min (Bandelin Sonorex R K512 H, Germany). The pH was adjusted to 9 with NH_4_OH, and the dispersion was kept under stirring at 180 °C overnight. Then, excess iron was removed with 5 centrifugation cycles at 4000 rpm for 15 min, and the final precipitate was washed by filtration using 5 L of water. Finally, the magnetized particles retained in the filter (rGNP@) were further washed by centrifugation as described above.

### Particle size and zeta potential

GNP, GNP-ox, and rGNP@ particle size distributions were determined with a LS230 laser particle analyzer (Coulter, United States). Data were collected performing 3 scans of 60 s, including polarization intensity differential scattering using Fraunhofer’s model. Zeta potential measurements were performed in a Zetasizer Nano ZS (Malvern Instruments, United Kingdom). Dispersions at 50 μg mL^-1^ were used for both particle size and zeta potential measurements.

### Preparation of PCL/rGNP@ composites

PCL/rGNP@ composites were prepared by melt-blending (Haake Polylab internal mixer, ThermoFisher Scientific, the Netherlands) with an internal mixing volume of 60 cm^3^, at a temperature of 90 °C, rotor speed of 200 rpm, and mixing time of 5 min. PCL/rGNP@ loads were of 2 and 10 wt.% rGNP@, for a total of 5 g per sample (PCL + rGNP@). The resulting composites were labeled PCL/rGNP@-2% and PCL/rGNP@-10%,respectively, and extruded as filaments with a maximum diameter of 3 mm.

### Chemical and thermal characterization

Fourier-transform infrared (FTIR) spectra were recorder using a Vertex 70 spectrometer (Bruker, Germany) in transmittance mode at 23 °C, coupled with an A225/Q Platinum diamond single-reflection accessory for attenuated total reflection (ATR). Spectra were recorded in the wavenumber range of 4000–400 cm^-1^ with an average of 60 scans and a resolution of 4 cm^-1^. Thermograms were recorded using a Polyma 214 differential scanning calorimeter (DSC; Netzsch, Germany) from samples weighing between 6 and 10 mg. Heating and cooling cycles were performed twice in a nitrogen gas atmosphere, heating from 23 to 150 °C at a rate of 10 °C min^-1^, followed by cooling to -20 °C at a rate of 10 °C min^-1^. The degree of crystallinity (*X*_*c*_) of PCL and PCL/rGNP@ composites was determined as follows:

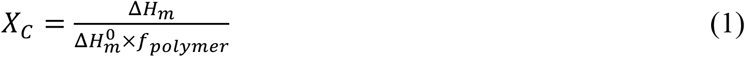

where *H*_*m*_ is the melting enthalpy or specific heat of fusion, *H*_*m0*_ is the melting enthalpy of 100% crystalline PCL, and *f*_*polymer*_ is the weight fraction of polymer in the sample. The *H*_*m0*_ of PCL was taken as 139.5 J g^-1 [33]^.

### Melt electrowriting

PCL and PCL/rGNP@ composites were melt-electrowritten (MEW) using an in-house built device. Briefly, PCL and PCL/rGNP@ composites were molten at temperatures (*T*) between 90 and 120 ºC in glass syringes with 27-G stainless steel nozzles (inner diameter = 0.2 mm) and pneumatically extruded at a pressure range of *P* = 1–3 bar (proportional pressure regulator, Festo, Germany), a positive applied voltage range of *V* = 5–7 kV (LNC 30000, Heinzinger Power Supplies, Germany), and a fixed collector distance of 4 mm. MEW jets were collected on a grounded aluminum collector plate driven by a triaxial motor controller (Trio Motion Technology Ltd., United Kingdom). PCL and PCL/rGNP@ fibers were printed at a range of collection speeds (*S* = 2–10 mm s^-1^), and the jets were monitored during printing with a Dino-Lite digital microscope (AnMo Electronics, Taiwan). The minimal speed at which the jet was deposited as a straight fiber was identified as the critical translation speed (CTS) for the corresponding set of *P, V*, and *T* values. The sets of PCL and PCL/rGNP@ fibers printed at different *P, V*, and *T* values were imaged with a stereomicroscope (Olympus), and fiber diameter were measured using ImageJ software (version 1.53; National Institutes of Health, United States). True (*d*_*F*_) and apparent fiber diameters (*d*_*F*_’) were quantified for straight and sinusoidal/coiled fibers, respectively. PCL and PCL/rGNP@ MEW scaffolds were designed with hexagonal pores with a side length of 0.6 mm and a scaffold thickness of 0.4 mm, and fabricated with the *P, V*, and *T* values that targeted fiber diameters of about 20 μm, as identified during diameter measurement of printed single fibers.

### Materials and printed scaffolds imaging

GNP powders, extruded filaments (cross-sections performed by cutting with a sharp blade), and MEW scaffolds were mounted on conductive carbon strips for visualization. Scanning electron microscopy and energy dispersive x-ray spectroscopy (SEM/EDS) analysis was performed using a FEI Quanta 400 FEG ESEM / EDAX Genesis X4M (ThermoFisher Scientific, USA) at an acceleration voltage of 3 kV. Particle diameter and aspect ratio of GNP, GNP-ox, and rGNP@ were determined by transmission electron microscopy (TEM). To prepare samples for imaging, aqueous dispersions of GNP, GNP-ox, and rGNP@ at a concentration of 50 μg mL^-1^ were sonicated for 1 h in a Sonorex R K512H ultrasound bath (Bandelin, Germany). Prior to imaging, dispersions were sonicated again for 10 min, and 10 μL of each were deposited on a carbon-coated TEM grid; after 1 min, excess material was removed using filter paper. TEM was performed at an acceleration voltage of 80 kV using a JEM 1400 microscope (JEOL, Japan) coupled with a CCD digital camera (Orious 1100 W; Hamamatsu Photonics, Japan). Images were processed using ImageJ.

### Printing accuracy assessment

SEM micrographs of MEW scaffolds were processed with threshold and segmentation analyzes using ImageJ software, and printed pore area was quantified for PCL, PCL/rGNP@-2%, and PCL/rGNP@-10% scaffolds. Accuracy of printed pores was assessed using the quality number (*Q*) as reported previously ^[34]^, defined as the ratio of printed pore area (*A*_*print*_) to expected pore area (*A*_*exp*_), as follows:

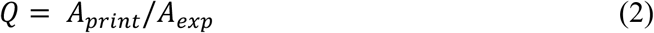

In case of astray printed fibers deposited across a pore, the *A*_*print*_ value is reduced, thus always leading to *A*_*print*_ < *A*_*exp*_, with a maximal value of *Q* = 1 corresponding to a perfectly accurate printed scaffold.

### Crystallographic and magnetic characterization

X-ray diffraction (XRD) analyses were obtained using a Smartlab diffractometer (Rigaku, United States) with a Bragg-Brentano geometry over a range of *2θ* = 15–70° at 23 °C and a Cu_Kα_ radiation beam (λ = 1.540593 Å). Magnetization (*M*) as a function of applied magnetic field (*H*) was measured using a commercial superconducting quantum interference device (SQUID) magnetometer (Quantum Design, Germany) at 310 K with a maximum magnetic field of 50 kOe.

### Tensile tests

PCL, PCL/rGNP@-2%, and PCL/rGNP@-10% composites were evaluated under monotonic uniaxial tension as extruded filaments (active length of 7 mm, diameter between 0.14 and 0.22 mm, *n* = 5) and MEW scaffolds (active area: width of 7 mm, length of 10 mm, thickness of 0.4 mm, pore size of 0.6 mm). Tests were performed using a 25-N force gauge in a Multitest 2.5-dV mechanical tester (Mecmesin, United Kingdom) at a strain rate of 1 mm min^-1^ at room temperature. MEW scaffolds were tested parallel to the main printing direction (x-direction, *n* = 3) and perpendicular to the main printing direction (y-direction, *n* = 3). Force-displacement curves were recorded using VectorPro software (Mecmesin), normalized to obtain engineering stress-strain curves, and processed to calculate the elastic modulus (for filaments), tangent modulus (for MEW scaffolds), yield strain, and elastic strain energy density (for filaments and MEW scaffolds). Elastic and tangent moduli were determined from least square fitting of the slope in the initial linear region of the engineering stress-strain curves for a coefficient of determination R^2^ > 0.99. Yield strain was established as the beginning of nonlinear deformation, thus indicated by the upper limit value of the range used for moduli calculations. Elastic strain energy density was determined as the integral of the engineering stress-strain curves from the origin to the yield strain, approximated numerically by the trapezoid method.

### Magnetic actuation experiments

PCL/rGNP@-2% and PCL/rGNP@-10% MEW scaffolds (thickness of 0.4 mm, pore size of 0.6 mm) were cut into strips (3 by 30 mm) and weighed. Apparent density (*ρ*_*app*_) was calculated from each MEW strip’s length (*L*), width (*w*), thickness (*t*), and weight (*m*) as follows:

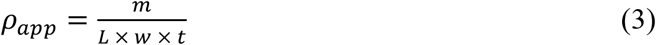

For testing, a strip was placed between an uncoated glass slide (1-mm thickness) and a custom-made support with a window (constant width of *x* = 15 mm), as shown in Figure 4E. The main plane of the strip was aligned to the gravitational force, so that its own weight did not alter the observed deflection. A NdFeB permanent magnet (100-kg strength; Supermagnete, the Netherlands) was placed at different distances from the strip, and the magnetic flux intensity was measured at the strip’s position with a magnetic field meter (MFM 3500; PCE Instruments, the Netherlands). Strip deflection was monitored with an EOS Rebel T3i DSLR camera (Canon Inc., Japan), and all tests were performed at room temperature. Magnetically triggered deflection with respect to magnetic flux intensity was measured using ImageJ software in terms of effective decrease in end-to-end length along the strip main axis (Δ*L/x*) and maximum protrusion from the main axis (*d/x*).

### Cell culture and construct assembly

The C2C12 cell line was obtained from ATCC (ATCC® CRL-1772™– Global Biosource Center, United States) and cultured in 10 cm dishes under standard conditions with Dulbecco’s Modified Eagle Medium (DMEM, Gibco™) supplemented with 20% fetal bovine serum (FBS; Biowest, batch S00F9). At 80% confluency, cells were harvested and counted to be used in the scaffolds. MEW scaffolds were cut into the desired shape using 6-mm biopsy punches. Prior to use, scaffolds were UV-sterilized for 15 min on each side, coated with extracellular matrix (ECM Gel from Engelbreth-Holm-Swarm murine sarcoma, Sigma) diluted at a 1:10 ratio in DMEM, and incubated for 30 min at 37 °C. Cells were encapsulated in a Matrigel (Matrigel® hESC-Qualified Matrix, LDEV-free; Corning, United States) / type-I collagen (Gibco™ collagen type I, rat tail, ThermoFisher) matrix with a 1:1:2 volume ratio. Matrigel was thawed on ice. Type-I collagen gel (2 mg mL^-1^) was prepared according to manufacturer instructions. Briefly, type-I collagen was mixed with 10x phosphate buffered saline (PBS) and DMEM and was neutralized with 1N NaOH. Encapsulation was done on ice. Cells and gels were mixed together thoroughly using a gel pipet. MEW scaffolds were placed in a Teflon mold between equal volumes of cell/gel mixtures and incubated for 30 min at 37°C to induce collagen gelation. The resulting constructs were cultured in 48-well plates (Corning) with 800 μL of culture medium consisting of DMEM with 20% v/v FBS. After 48 h, culture medium was replaced by differentiation medium consisting in DMEM and 2% v/v horse serum (Gibco™) and samples were cultured for 6 more days without medium replacement. For 2D differentiation, C2C12 cells were cultured under normal conditions on μ-slides (Ibidi, Germany) coated with ECM (1:10 ratio in DMEM). At 80% confluency, culture medium was replaced for differentiation medium and cultured for 6 more days.

### Live/dead staining and quantification

ReadyProbes™ Cell Viability Imaging Kit, Blue/Green (ThermoFisher, cat. R37609) was used to perform live/dead staining. Briefly, two drops of NucBlue® Live reagent (Hoechst 33342) and two drops of NucGreen® Dead reagent were added to 1 mL of medium. Samples were then incubated for 30 min at room temperature, followed by fluorescence imaging. NucBlue® Live reagent was used to stain nuclei of all cells and detected with a standard 4′,6-diamidino-2-phenylindole (DAPI, blue) filter. NucGreen® Dead reagent was used to stain only the nuclei of cells with compromised plasma membrane integrity and was detected using a standard FITC (green) filter. Fiji/ImageJ software (Version 2.0.0-rc-69/1.52p) was used to quantify the total amount of cells and the total amount of dead cells. Briefly, blue and green channels in the images were separated and converted to 8-bit. Next, the function ‘Find Maxima’ was used to identify the cells. The output was used to calculate the percentage of live cells.

### Fluorescence staining and confocal imaging

Samples were fixed using 2% v/v paraformaldehyde for 20 min at room temperature following three rinsing steps in PBS. Samples were permeabilized and blocked in PBS, 0.1% v/v Triton-X-100, and 5% v/v goat serum for 60 min. Samples were then incubated with the primary antibody MF-20 (DSHB Biology) at a concentration of 1:60 in 5% v/v goat serum at 4 °C overnight followed by three rinsing steps in PBS. Next, samples were incubated with Alexa Fluor® 488 Goat anti-Mouse IgG (H+L) (Life Technologies, cat. A11001) in PBS at a concentration of 1:500 for 60 min at room temperature. Finally, samples were rinsed three times with PBS of which the second step is with DAPI at 1:5000 (Life Technologies, cat. D1306). For imaging, microscopy slides with Press-To-Seal Silicone Isolators (Bio-Labs-JTR8S-1.0) were used. Samples were embedded in mounting media (50% v/v glycerol in PBS) and imaged on a Dragonfly200 confocal microscope. Full scaffolds were imaged and processed for analysis.

### Quantification of cell fusion and differentiation

Images from the whole scaffolds (acquired using Dragonfly200) were quantified using a pipeline developed on open-source CellProfiler software ^[35]^. Briefly, nuclear staining and myotube staining images were loaded individually into the same pipeline. For nuclei identification, pre-processing steps were applied to all images to enhance image features and a median filter to reduce unspecific object identification. For myotube identification, a Gaussian filter was applied to the images before object identification was used to measure object morphology. Two filter steps were applied to myotube images to reduce non-specific segmentation and Otsu thresholding method was used to separate myotubes. A mask was applied to capture the nuclei within the myotubes, and the outputs were related to each other. To assess the efficiency of differentiation, two parameters were evaluated. Undifferentiated myoblasts contain one nucleus, and myotubes contain multiple nuclei. First, the fusion index (FI) was calculated as the number of total nuclei within the sample divided by the number of nuclei within myotubes. To further characterize differentiation, the number of nuclei per individual myotube was determined and identified as the myoblast/myotube ratio.

### Statistics

All statistical analysis was carried out using Prism 8 software (GraphPad Software, United States). Statistical significance between groups was assessed as indicated in each case. Differences were considered significant for p < 0.05 and labeled as * for p < 0.05, ** for p < 0.01, *** for p < 0.001, and **** for p < 0.0001. Data are plotted as mean ± standard deviation.

## Results and discussion

### 1. Synthesis of magnetized particles and magnetic PCL composites

In this study, a simultaneous reduction and magnetization of GNP-ox was successfully performed to yield rGNP/ION (labeled as rGNP@). This powder was then melt-blended with medical-grade PCL to yield highly magnetic PCL/rGNP@ composites (Figure 1). First, GNP were oxidized by the modified Hummers method (MHM) to yield GNP-ox. Then, a redox reaction involving FeCl_2_ and GNP-ox allowed for ION deposition *in situ* onto self-reduced graphene nanoplatelet surface to yield rGNP@. The reaction was successful, resulting in well-dispersed ION at rGNP surface (Figure 1A,B). GNP, GNP-ox, and rGNP@ powders presented distributions of particle size in the microscale, almost entirely below 2 μm for the three powders (Figure 1C). In particular, rGNP@ showed a mean hydrodynamic particle size of 0.96 ± 0.50 μm, which was slightly greater than for GNP-ox (0.89 ± 0.34 μm, Figure S1A), thus indicating that hydrodynamic particle size did not significantly increase after ION deposition. Moreover, the zeta potential of rGNP@ is lower than that of non-deposited ION, indicating that the former have higher water stability (Figure S1B). Such features were preferred to ensure biocompatibility, since poor water dispersability has been associated with a lack of biodegradation, bioaccumulation, and toxicity of nanomaterials ^[36,37]^, while large particles (above 200 nm) are more often phagocytosed than internalized ^[38]^.

**Figure 1.**
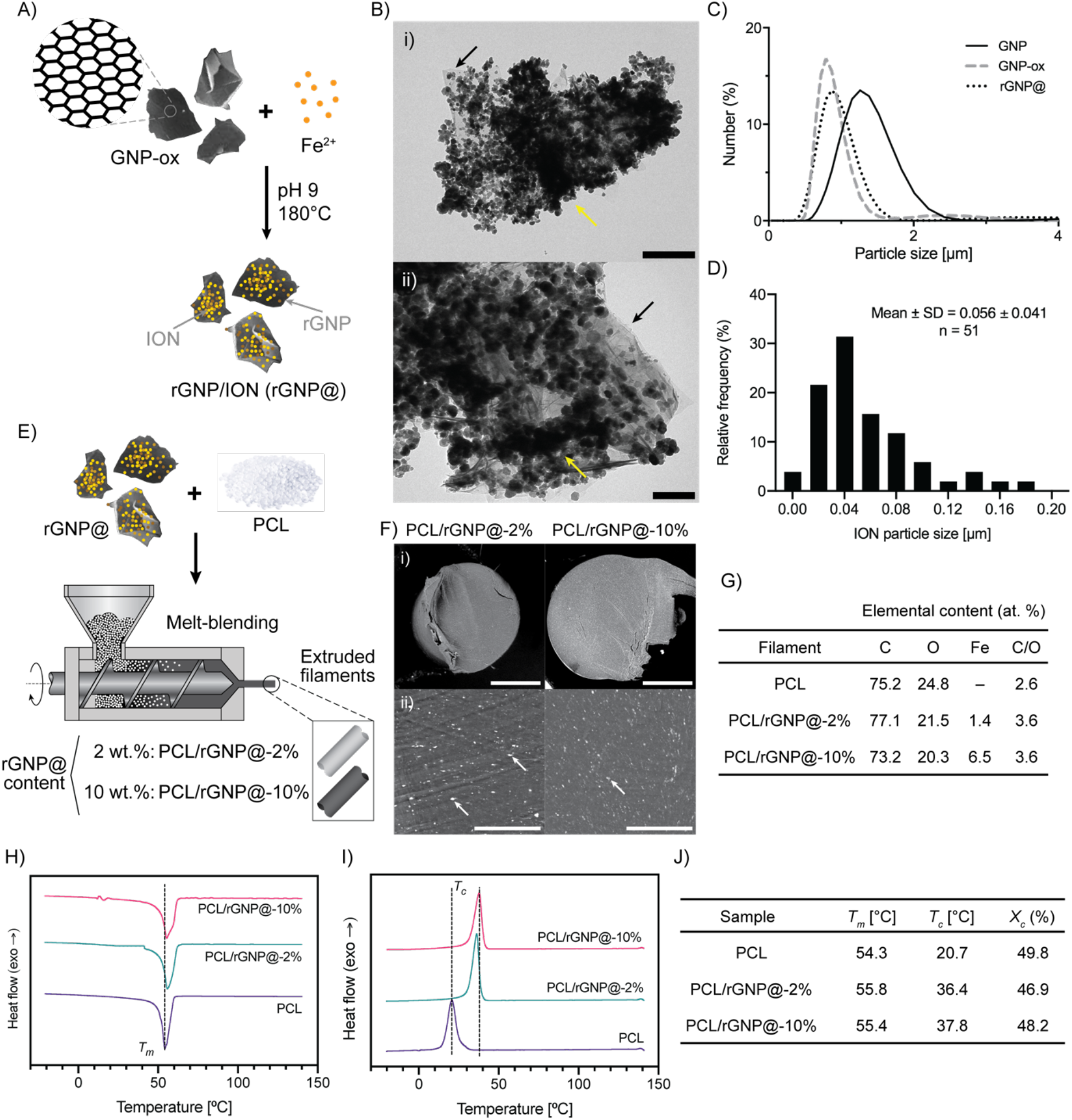
Production of magnetic particles and magnetic PCL composites. A) Schematic of synthesis of magnetized graphene nanoplatelets (rGNP@) by deposition of iron oxide nanoparticles (ION) on oxidized graphene nanoplatelets (GNP-ox) *in situ*. B) TEM images of rGNP@ particles; yellow arrows indicate ION, and black arrows indicate GNP-ox particles. Scale bars: i) 500 nm, ii) 200 nm. C) Particle size distribution of graphene nanoplatelets (G), GNP-ox, and rGNP@ as determined by DLS; polydispersity indices (PDI) for each individual particle population are shown. D) ION particle size distribution as determined from TEM. E) Schematic of preparation of magnetic composites from poly-(ε-caprolactone) (PCL) and rGNP@ at 2, and 10 wt%. F) SEM images of PCL/rGNP@-2% and PCL/rGNP@-10% filaments, showing the distribution of rGNP@ particles (white dots indicated by white arrows) in the PCL matrix (gray); scale bars: i) 500 μm, ii) 50 μm. G) Elemental composition of PCL/rGNP@ composites from EDS analysis. H) DSC second heating and I) second cooling cycle curves of PCL and PCL/rGNP@ composites, showing peak melting and peak crystallization temperatures (*T*_*m*_, *T*_*c*_). J) Transition temperature and degree of crystallinity (*X*_*c*_) data of PCL and PCL/rGNP@ composites, as obtained from DSC.

Furthermore, particle morphology has been identified as a mechanism for ION cytotoxicity. For instance, rod-shaped ION with aspect ratios between 5 and 10 have been found to undergo nonspecific cellular internalization, leading to heightened inflammatory responses in macrophages compared to spherical ION, potentially due to the greater contact area, sharper features, and membrane penetrating capability of elongated particles ^[39]^. Therefore, in addition to analyzing hydrodynamic particle sizes, we investigated the particle size, aspect ratio, and shape of GNP, GNP-ox, and rGNP@ powders by TEM. Individual ION deposited on GNP were observed to have a wide monomodal particle size distribution (mean = 56 ± 41 nm; Figure 1D) and an aspect ratio ranged from 1 to 2, while GNP, GNP-ox, and rGNP@ particle diameters ranged from 0.5 to 2 μm (Figure S1C–D). Individual GNP can only be observed under TEM after oxidation (GNP-ox); however, the magnetization process covers most of the graphene surface area with generally globular ION (Figure S1E). In addition, microscale investigation of elemental composition and spatial distribution by SEM-EDS revealed that rGNP@ contains a homogeneous distribution of iron, amounting to a total iron content of 37 atomic % (= 69 weight %; Figure S1F–G), thus confirming the potential of rGNP@ powder as a filler to introduce magnetic properties into polymer composites.

PCL is a thermoplastic and the gold standard material for MEW of synthetic scaffolds, due to its biodegradability, high biocompatibility, low melting point (56–60 °C), and stable thermal and rheological properties which facilitate processing in the molten state for hours ^[40]^. Thus, rGNP@ were incorporated into PCL by melt-blending in order to impart magnetic properties to the polymer, while permitting controlled MEW processing of the composite due to low rGNP@ agglomeration (Figure 1E). Melt-blending at 90 °C yielded filaments of PCL compounded with 2 and 10 wt.% rGNP@ particles (PCL/rGNP@-2% and PCL/rGNP@-10%, respectively, Figure S2). Under SEM, rGNP@ were observed in the composites throughout the entire filament cross-sections as well-dispersed bright speckles due to the high conductivity of reduced GNP (Figure 1F). In addition, elemental analysis confirmed successful loading of ION within the PCL matrix (Figure 1G), while FTIR analysis confirmed the incorporation of rGNP@, as indicated by the rGNP@ absorption band present in the PCL/rGNP@ composites (broad band at 400–700 cm^-1^; Figure S2E). The addition of rGNP@ only slightly increased the composite melting point (T_M_) with respect to pristine PCL; however, the crystallization temperature (T_C_) increased considerably in proportion with rGNP@ content, from 20.7 °C for PCL to up to 37.8 °C for PCL/rGNP@-10% (Figure 1H–J). This indicates that rGNP@ increases the crystallization rate of pristine PCL in a concentration-dependent monotonic manner, suggesting that rGNP@ acts as a crystallization nucleating agent in the PCL matrix. Nevertheless, PCL with rGNP@ loads of up to 10 wt.% were found to remain extrudable (Figure 2). Also, a content of 10 wt.% rGNP@ represents the highest content of graphene-based fillers embedded in PCL matrices that has been processed by MEW and reported thus far in the literature. Previous reports have demonstrated MEW of PCL composites obtained by solution mixing, which is an approach limited to yielding composites with low particle content, including reduced graphene oxide content of up to 1 wt.% ^[28]^ and ION up to 0.3 wt.% ^[30]^. Meanwhile, melt-blending has been reported to produce PCL/graphene composites with up to 9 wt.% graphene, processable by conventional fused deposition manufacturing, and able to produce filaments with diameter above 0.3 mm ^[37,41]^, but these composites have not yet been implemented in the context of microscale fibers, *i*.*e*. fibers with diameter below 0.3 mm. However, in this work we report for the first time that melt-blending allows for the successful incorporation of a very high content of rGNP/ION (up to 10 wt.%) in PCL matrices, yielding highly homogeneous composites with appropriate processability for MEW.

**Figure 2.**
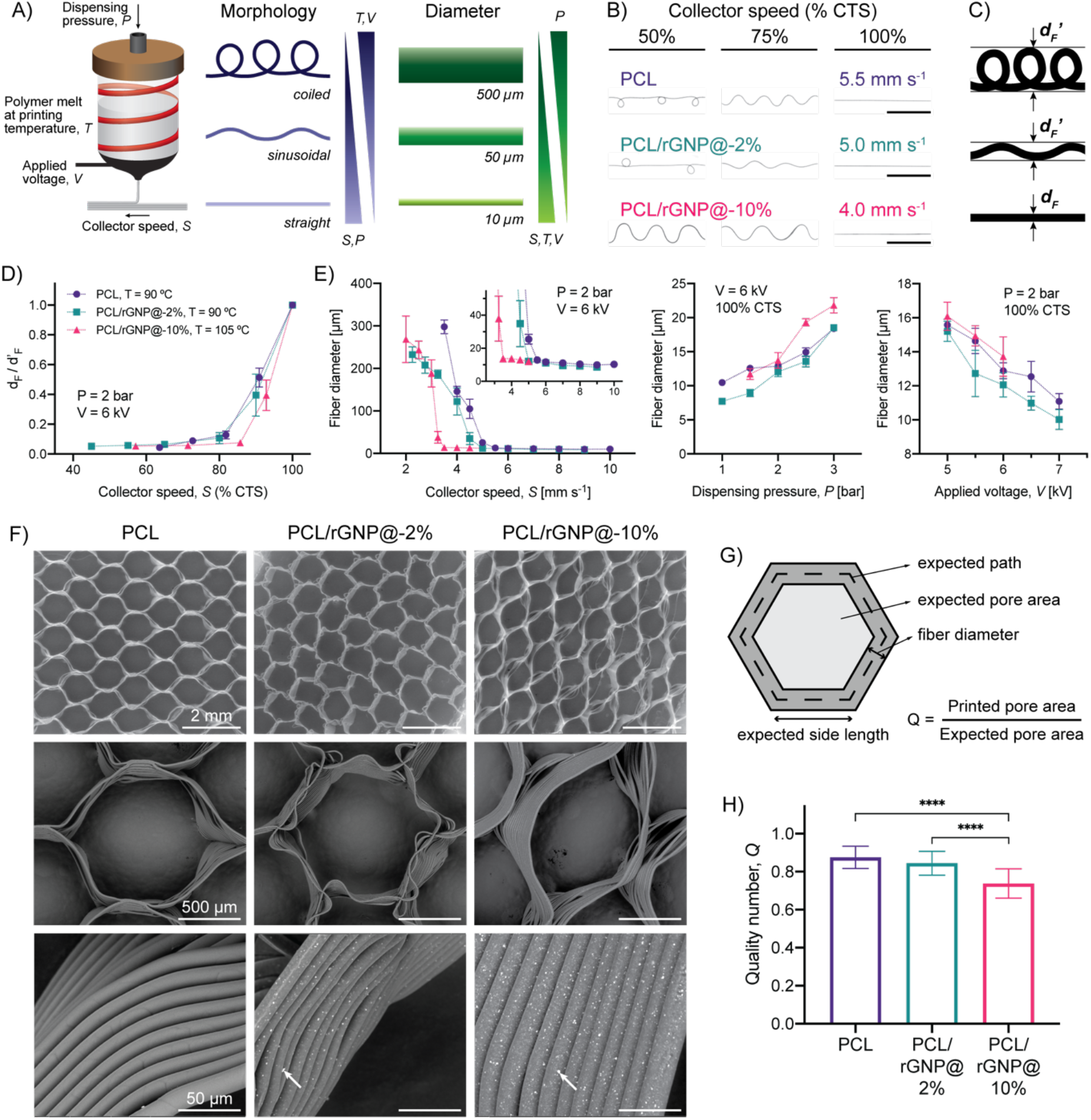
MEW of magnetic PCL composites into fibers and assembly into 3D hexagonal scaffolds. A) Representation of the effect of MEW printing parameters on fiber morphology and diameter. B) Representative micrographs of MEW fibers showing the dependence of morphology on composition and collector speed; the collector speed identified as critical translation speed (CTS) is shown. Scale bars: 1 mm. C) Schematic showing the definition of observed fiber diameter in straight (true diameter *d*_*F*_), sinusoidal, or coiled fibers (apparent diameter *d*_*F*_’). D) True-to-apparent diameter ratio (*d*_*F*_*/d*_*F*_’) between straight and coiled or sinusoidal fibers, indicating the dependence of morphology on collector speed. E) Effect of composition, collector speed, dispensing pressure, and applied voltage on MEW fiber diameter. F) SEM micrographs of PCL and PCL/rGNP@ composite MEW scaffolds (rGNP@ particles indicated by white arrows). G) Schematic of the expected features in the MEW scaffold design and the definition of the quality number, *Q*, as indicator of print fidelity. H) Effect of rGNP@ content on the quality number of MEW scaffolds (Kruskal-Wallis test with Dunn’s multiple comparisons).

### 2. MEW of magnetic PCL/rGNP@ composites and integration into multi-material well-organized fibrous scaffolds

A wide range of printing parameters determine the morphology and diameter of a MEW fiber. Here, we assessed the effect of key instrument parameters, *i*.*e*. dispensing pressure (*P*), printing temperature (*T*), applied voltage (*V*), and collector speed (*S*) on the morphology and diameter of PCL and PCL/rGNP@ single fibers (Figure 2A). Given a fixed set of *P, T*, and *V* parameters, *S* was varied to determine the lowest value at which the printed fiber has a straight morphology, thus identified as the critical translation speed (CTS), which generally decreased with increasing rGNP@ content (Figures 2B and S3), in agreement with the greater loss factor of high-rGNP@ composites (Figure 1J). To quantitatively assess the influence of printing parameters on fiber morphology, the true-to-apparent diameter ratio (*d*_*F*_*/d*_*F*_’) was measured for single MEW fibers, considering a value of *d*_*F*_*/d*_*F*_’ = 1 for a perfectly straight fiber (Figure 2C). Due to the significant rheological differences across PCL/rGNP@ composites, fibers were printed at different *T* in order to tune *P* and *V* values within the operational range of the MEW device (0–6 bar and 1–10 kV, respectively); PCL and PCL/rGNP@-2% were printed at 90 °C and PCL/rGNP@-10% at 105 °C. With fixed values of *P* (= 2 bar) and *V* (= 6 kV) for all compositions, *S* values were normalized as a fraction of the CTS; this allowed to observe a consistent trend in *d*_*F*_*/d*_*F*_’ values regardless of composition. Values of d_F_/d_F_’ showed an asymptotic-like increase with S at low collector speeds and a sharp increase at *S* above 85% CTS (Figure 2D). Importantly, *d*_*F*_*/d*_*F*_’ values showed high variability (relative error below 25%) only in the *S* range of 85–100% CTS (Figure 2D), which is ascribed to the variability of MEW fibers with sinusoidal morphology close to straight morphology, since small local heterogeneities in jet temperature and viscosity can severely alter the amplitude of sinusoidal fibers. Additionally, we assessed the effect of *S, P*, and *V* on fiber diameter; when a parameter was varied, the others were kept at *P* = 2 bar, *V* = 6 kV, and *S* = 100% CTS (Figure 2E). At low *S* values under the CTS, the apparent fiber diameter decreases with increasing speed due to the coiled-to-straight transition. Meanwhile, above the CTS, fibers of all compositions have straight morphology, and at high *S* values, diameters approach the range of 8–13 μm with relatively low dependence on *S*, as reported previously ^[42]^. Overall, for PCL, PCL/rGNP@-2%, and PCL/rGNP@-10% samples, *P* and *V* can be adjusted to modulate MEW fiber diameter, for instance, from 8 to 20 μm (Figure 2E).

PCL and PCL/rGNP@ composites were successfully processed into MEW scaffolds with a hexagonal microstructure (Figure 2F–H). This hexagonal geometry was selected since we have previously described to display elastic deformability under tensile loading and scaffold moduli values about 50 to 80% lower than for rudimentary linear microstructures, generally with square or rectangle geometries ^[10]^, thus facilitating deformation under low forces, as is the case in remote stimulation of *in vitro* tissue models. Moreover, conventional square, or rectangular, crosshatch microstructures fail at relatively low strains (∼5%) ^[15]^ and therefore are not compatible with physiological strains experienced by skeletal muscle. For these overall reasons, linear microstructures like square or rectangular MEW geometries were not considered for this study. SEM imaging revealed that MEW of PCL and PCL/rGNP@-2% enabled more accurate fiber stacking in scaffolds with thickness up to 0.4 mm and hexagonal side length of 0.6 mm, whereas PCL/rGNP@-10% presented greater pore size variation and higher incidence of fiber misplacement and fiber wall slanting (Figure 2F). These printing defects in high-rGNP@ composites cumulatively reduce the average pore size and pore homogeneity. To quantitatively evaluate the printing accuracy of PCL composites with different rGNP@ content, the quality number *Q* was defined as the ratio between printed and expected pore area (Figure 2G), as reported previously for similar electrowritten fiber scaffolds ^[34]^. In agreement with SEM observations, printing accuracy decreased with increasing rGNP@ content, and *Q* values were lower for PCL/rGNP@-10% (mean of 0.74) than for PCL and PCL/rGNP@-2%, and PCL/rGNP@-10% (mean of 0.88 and 0.84, respectively; Figure 2H). This can be ascribed to the decreasing processability of composites with increasing rGNP@ content above 10 wt.%. Despite this limitation, MEW provided an overall robust approach for processing composites with high filler content up to 10% into scaffolds with controlled organization and geometry.

In order to assess whether the MEW process altered the magnetic and crystallographic properties of PCL/rGNP@ composites, magnetometry and XRD studies were performed on GNP powders and melt-extruded filaments and compared to MEW scaffolds (Figure 3, S4). The H-M hysteresis curves of GNP, GNP-ox, and rGNP@ were in agreement with typical ferromagnetic behavior (Figure S4A). Importantly, rGNP@ showed a high saturation magnetization (*M*_*S*_) value of 79.2 emu g^-1^ despite the presence of GNP, which has very low *M*_*S*_ (Figure S4B). The magnetization of rGNP@ was found to be superior to those of other reported GBM/magnetic fillers, such as ION-decorated carbon nanotubes, with *M*_*S*_ ranging from 25 to 40 emu g^-1 [43]^. In contrast to carbon nanotubes, GNP have a larger surface area that increases the amount of deposited ION. In addition, melt-blended composite filaments showed *H*-*M* curves with typical ferromagnetic behavior and *M*_*S*_ values of 1.4 and 4.6 emu g^-1^ for PCL/rGNP@-2% and PCL/rGNP@-10%, respectively, while the MEW scaffolds showed *M*_*S*_ values of 1.2 and 4.5 emu g^-1^ for PCL/rGNP@-2%, and PCL/rGNP@-10%, respectively (Figure 3A,D). These *M*_*S*_ values are comparable to those of previously reported PCL/ION composites lacking graphene-based materials, which range from 1 to 3 emu g^-1^ for filler contents from 5 to 10 wt.% ^[44]^. Moreover, XRD analysis confirmed the oxidation of GNP into GNP-ox, since in the GNP-ox pattern the characteristic (002) reflection peak of graphite disappears (*2θ* = 26.5°) and a broad (001) peak ascribed to graphene oxide emerges (centered at *2θ* = 12°; Figure S4C) ^[45]^. The absence of this broad graphene oxide peak in the rGNP@ pattern confirms the partial reduction of GNP-ox during the ION deposition reaction (Figure 3B). Additionally, the XRD reflection peaks of rGNP@ were associated to the cubic spinel structure with *Fd3m* symmetry, typical of ferrites (Figure 3B). Moreover, rGNP@ presented a calculated lattice parameter of *a* = 8.364 Å, which lies between the parameters of magnetite Fe_3_O_4_ (*a* = 8.396 Å, JCPDS #19-0629) ^[46]^ and maghemite Fe_2_O_3_ (*a* = 8.346 Å, JCPDS #39-1346) ^[47]^, thus suggesting that the ION in rGNP@ consist of a mixture of both ferrites. The PCL pattern presented the characteristic peaks of the orthorhombic (110) and (200) crystalline planes at *2θ* = 21.8 and 24.2°, respectively ^[48]^. Overall, the XRD patterns of the PCL/rGNP@ composites showed the overlaying peaks of PCL and rGNP@, with their intensities varying in relation to rGNP@ content as expected and no significant shifts, for both as-extruded and MEW composites (Figure 3B,E). The *M*_*S*_ ratios between rGO@ and PCL/rGO@ and the XRD peak intensity ratios between PCL and rGO@ were related to the nominal rGO@ content of composites, showing similar trends and values for both as-extruded and MEW composites (Figure 3C,F). This confirms that the thermal and electrical conditions involved in the MEW process do not cause substantial decreases in magnetization or changes in crystalline phases of the ION components, thus validating MEW as a microfiber fabrication technique that preserves the magnetic properties of magneto-active thermoplastic composites.

**Figure 3.**
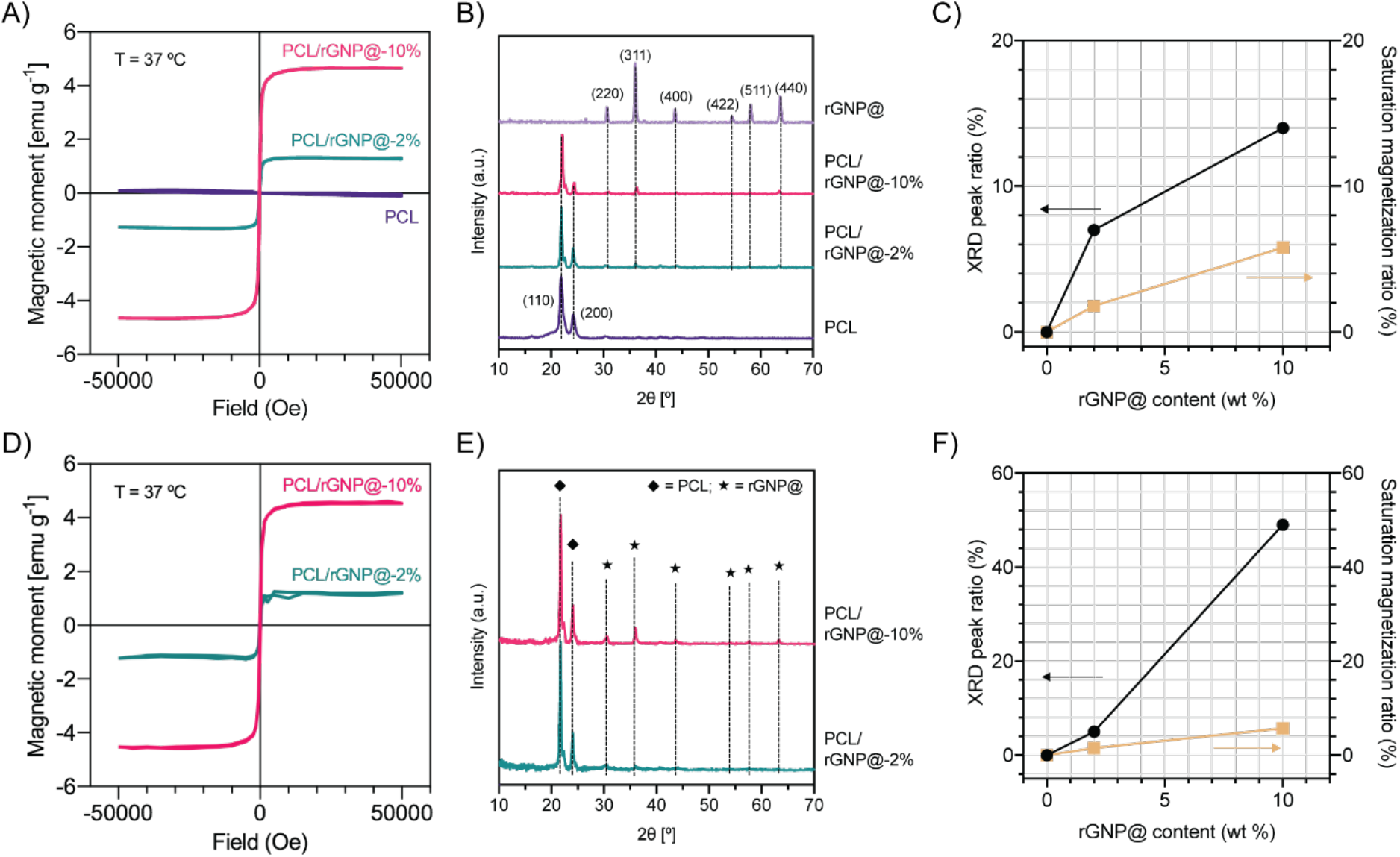
Crystallographic and magnetic properties of melt-extruded filaments and melt-electrowritten scaffolds. A) Magnetic hysteresis curves, B) x-ray diffractograms, and C) correlation between magnetic and crystallographic parameters of melt-blended PCL and PCL/rGNP@ filaments. D) Magnetic hysteresis curves, E) x-ray diffractograms, and F) correlation between magnetic and crystallographic parameters of PCL and PCL/rGNP@ melt-electrowritten scaffolds.

### 3. Mechanical behavior and magnetic actuation of PCL and PCL/rGNP@ MEW scaffolds

Uniaxial tensile testing of melt-extruded filaments with diameters between 0.14 and 0.22 mm was performed to evaluate the influence of rGNP@ incorporation and the extrusion process on PCL bulk material properties (Figure 4A–B). PCL/rGNP@-2% and PCL/rGNP@-10% filaments showed significantly higher elastic moduli, 357 ± 16 and 369 ± 20 MPa, respectively, when compared to PCL filaments, 316 ± 14 MPa (Figure 4B). This increase in the elastic modulus can be attributed to the filler incorporation in good agreement with previous studies. For instance, rGO has been reported to increase the elastic modulus of PCL composites with rGO content of 0.1 wt. %, both for melt-cast and melt-electrowritten composite fibers, when compared to pristine PCL ^[28]^. Similarly, ION incorporation increases the elastic modulus of PCL matrices, as observed in electrospun composite fibers ^[49]^. In general, stif particles such as ceramics and metals, when included in a PCL matrix, can act as anchor points that hinder the reversible PCL chain sliding that leads to elastic strain. In addition, the elastic modulus of melt-extruded pristine PCL filaments was observed to be considerably higher than that of melt-cast PCL with a similar molecular weight (about 130 MPa), as reported by Somszor *et al*. ^[28]^. This observation is expected, since all extrusion processes —including melt-blending during rGNP@ incorporation and printing— cause shear-induced alignment of the PCL chains and overall strengthening of the polymer matrix. Interestingly, no significant difference in elastic modulus, yield strain, and elastic strain energy density was observed among filaments with 2% and 10% rGNP@. This can potentially be explained by the fact that both rGNP@ concentrations allowed for uniform dispersion and interaction with the PCL matrix, without the introduction of defects. Moreover, there was no significant difference among the yield strain and elastic strain energy density values of all compositions tested (Figure 4B), showing that the incorporation of rGNP@ into PCL does not significantly alter the range of the elastic regime and only slightly increases the material’s intrinsic stifness.

**Figure 4.**
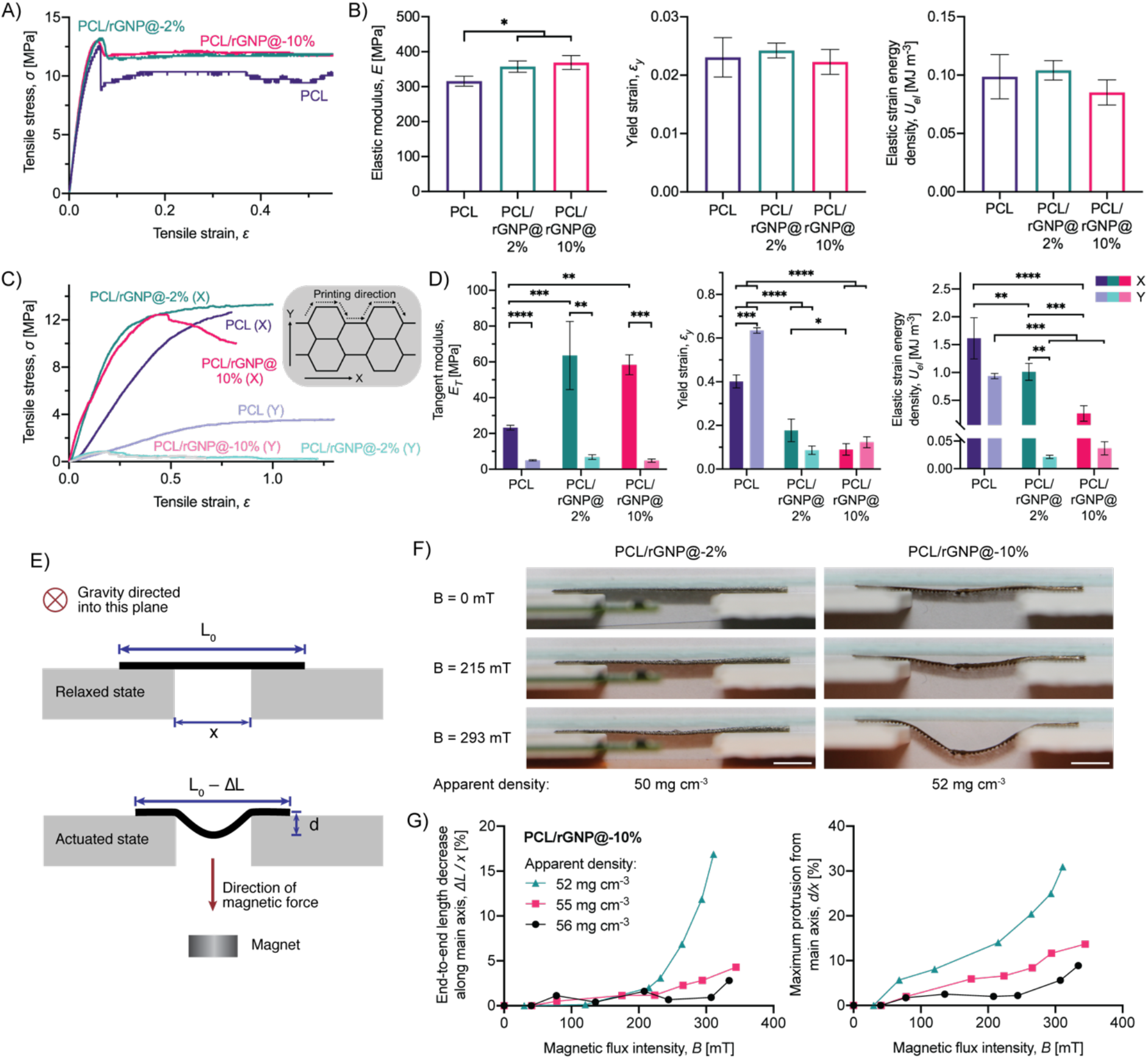
Mechanical behavior and actuation of magnetically deformable MEW scaffolds. A) Representative uniaxial stress-strain curves and B) mechanical parameters of PCL and PCL/rGNP@ extruded filaments (Kruskal-Wallis test with Dunn’s multiple comparisons). C) Representative uniaxial stress-strain curves and D) mechanical parameters of PCL and PCL/rGNP@ MEW scaffolds (two-way ANOVA with Sidak’s multiple comparisons test for groups with same direction; multiple unpaired t tests with Holm-Sidak multiple comparison correction for groups with same composition). E) Setup for assessing magnetic actuation on PCL/rGNP@ MEW scaffolds. F) Snapshots from magnetic actuation experiments on PCL/rGNP@ scaffolds; scale bars: 5 mm. G) Magnetic flux intensity (*B*)-deformation curves for PCL/rGNP@-10% scaffolds: end-to-end length decrease along the main axis (left), and protrusion away from the main axis (right).

The microfiber fabrication process via MEW introduced composition-dependent variations in scaffold mechanics that are not observed in the melt-extruded materials themselves (Figure 4C–D). In general, across all compositions, the values of tangent modulus, yield strain, and elastic strain energy density were significantly greater for scaffolds tested along the main printing direction (labeled as x-direction) than in the perpendicular direction (labeled as y-direction; Figure 4C–D), which is ascribed to two main reasons. First, the orientation of the printed hexagon edges determines that the x-direction is completely aligned to one pair of edges and partially aligned to two pairs, while the y-direction is only partially aligned to two pairs. Second, fibers are deposited continuously along the x-direction, whereas fusion between newly deposited and solidified fibers is achieved only partially, as can be observed from SEM micrographs (Figure 2F). Altogether, this leads to a greater degree of fiber bending and more potential for fiber delamination under tension along the y-direction. These observations are in good agreement with previous works on melt-electrowritten PCL hexagonal scaffolds ^[10]^. Regarding the effect of composition, the incorporation of rGNP@ significantly increases the tangent modulus values of 2% and 10% filler-containing scaffolds with respect to PCL along the x-direction, while there is no such significant change along the y-direction (Figure 4D). Additionally, the yield strain and elastic energy density are significantly reduced for composite scaffolds with respect to PCL scaffolds, indicating a greater component of irreversible deformation for composite scaffolds (Figure 4D), as observed previously in polymer/graphene composites with very high filler content ^[31,37]^. Overall, this can be explained considering that, in composite fibers, deformation causes the movement and reorientation of filler particles and PCL chain sliding around particles, both of which lead to a permanent loss of elastic energy. Nevertheless, the elastic behavior of the 2% and 10% filler-containing scaffolds was within the range observed in skeletal muscle, for instance, in the diaphragm of a mouse model of early-onset muscular dystrophy with myositis ^[50]^ and the tibialis anterior muscle of healthy adult Wistar rats ^[51]^.

To assess the capability for magnetically triggered deformation, composite MEW scaffolds were subjected to different magnetic field intensities in unconstrained conditions. The setup allowed to monitor scaffold out-of-plane bending through a window of constant width (*x*), which led to an effective decrease in end-to-end length along the scaffold main axis (Δ*L/x*) and maximum protrusion away from the main axis (*d/x*) in the actuated state (Figure 4E). Even at a magnetic flux intensity *B* = 293 mT, PCL/rGNP@-2% scaffolds did not perceive sufficient magnetic force to undergo observable bending; however, magnetic forces in PCL/rGNP@-10% scaffolds at *B* = 293 mT were high enough (Figure 4F). For this composition, magnetically triggered deformation was reversible and was found to increase gradually with the magnetic flux intensity (Figure S5). Importantly, a dependence of scaffold deformation on fiber density was observed; to assess this, scaffolds with different fiber densities, and thus different apparent densities, were tested. Both the effective main-axis length decrease and maximum protrusion values were observed to increase more notably for scaffolds with lower apparent density (Figure 4G), while at higher apparent density, scaffolds possess greater out-of-plane stifness that restrains bending. Greater deformation was found for PCL/rGNP@-10% scaffolds with apparent density of 52 mg cm^-3^, showing effective main-axis length decrease of 17% and maximum protrusion of 31% at *B* = 293 mT (Figure 4G). Interestingly, reversible actuation of PCL/rGNP@-10% scaffolds required only a magnet with 100-kg strength and maximum magnetic field intensity of about 300 mT. On the other hand, previously reported magneto-active scaffolds are often based on hydrogels or ceramics and require much higher magnetic field intensities to achieve deformation. For example, a hydroxyapatite sponge was reported to undergo contraction of up to 25% in close proximity to an electromagnet with 200-kg strength ^[25]^. Additionally, Spangenberg *et al*. reported that 3D-printed reticulated scaffolds based on ION-loaded alginate/methyl cellulose hydrogels were capable of contraction up to 4%, which occurred due to filament collapse under a magnetic field intensity of up to 200 mT ^[52]^. These hydrogel and ceramic scaffolds cannot undergo large magnetically triggered deformations, as they lack the wide elastic range of PCL and other thermoplastics, which can act as a reinforcing platform for stimulation. Although the development of thermoplastic/ION composites for cellular scaffolds has been addressed previously ^[30,44,53–55]^, efforts have focused on magnetic hyperthermia, imaging, and other applications of ION, without proper evaluation of the actuation potential of these composites. In the face of this issue, the results here presented confirm the potential of PCL/rGNP@ MEW scaffolds as tunable platforms for controlled out-of-plane actuation and magnetically induced mechanical stimulation.

### 4. Biological evaluation of PCL and PCL/rGNP@ MEW scaffolds

The biological potential of magnetic PCL for skeletal muscle engineering was assessed using C2C12 myoblasts. First, C2C12 cells were cultured on PCL scaffolds to identify an appropriate cell density. A cell number titration experiment was performed in which three starting concentrations of cells (50000, 75000, and 100000 per scaffold) were assessed. All three cell concentrations show cellular alignment along the scaffold fibers (Figure 5A). As a measure of differentiation efficiency, the fusion index was calculated. No significant difference of fusion index was observed between the three conditions (Figure 5B). However, a starting concentration of 100000 cells per scaffold showed a significant difference in differentiation efficiency, *i*.*e*. myoblast-to-myotube fusion, indicating that longer myotubes were formed. Therefore, this concentration was used in the remainder of the study.

**Figure 5.**
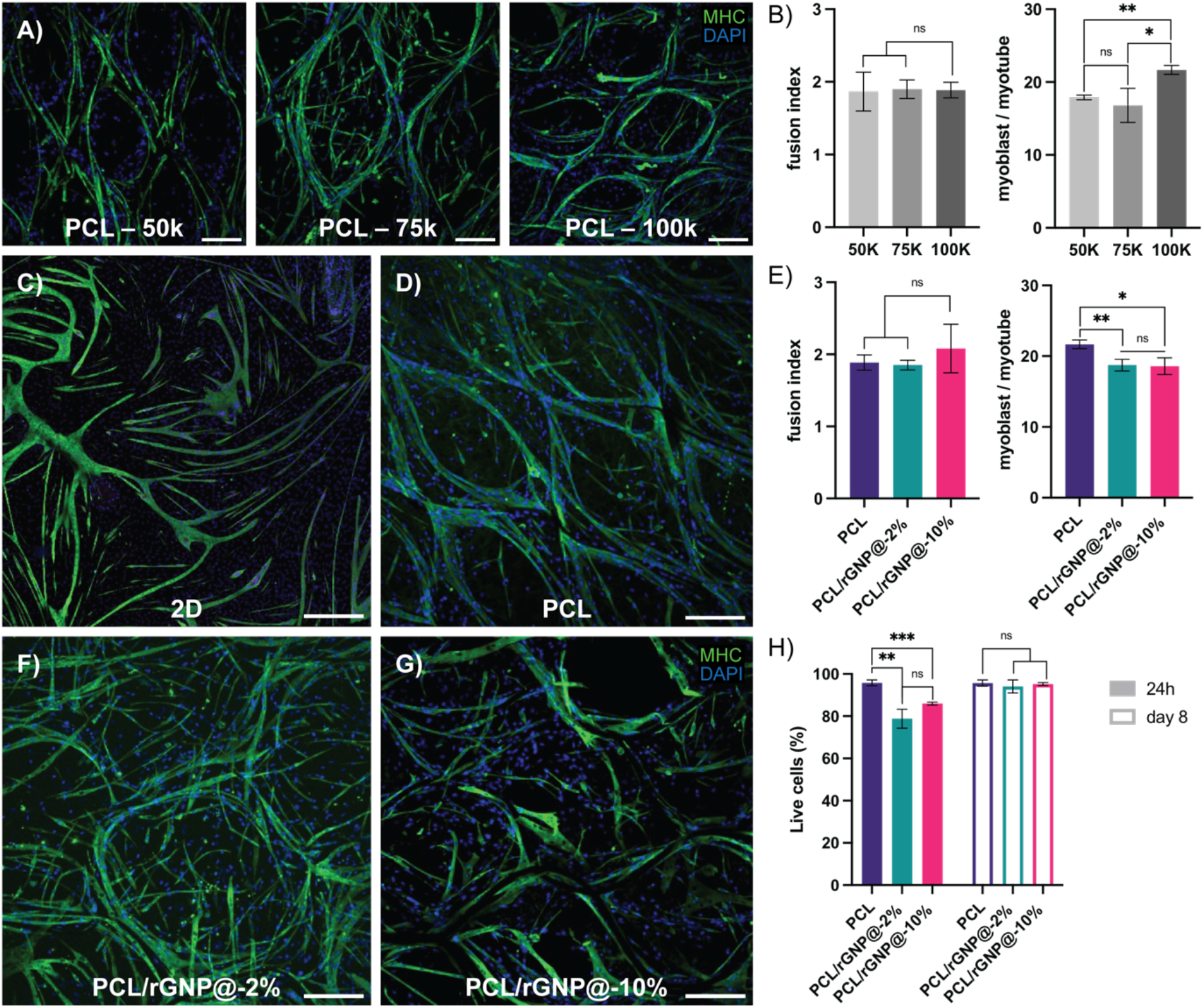
Biological evaluation of MEW PCL and PCL/rGNP@ scaffolds in Matrigel/collagen hydrogels. A) Confocal images of C2C12 in PCL scaffolds at three different cell numbers. B) Fusion index (determined as the ratio of total nuclei to nuclei in myotubes) and differentiation efficiency (myoblast/myotube, determined as the number of nuclei per myotube) at three different cell numbers (*n* = 3) (one-way ANOVA with Tukey multiple comparisons). C) Confocal image of C2C12 cells cultured in 2D conditions. D) Confocal image of C2C12 cells cultured in PCL scaffold. E) Fusion index and differentiation efficiency (myoblast/fiber) for PCL, PCL/rGNP@-2% and PCL/rGNP@-10% scaffolds (*n* = 3) (one-way ANOVA with Tukey multiple comparisons). F) Confocal image of C2C12 cells in PCL/rGNP@-2% scaffolds. G) Confocal image of C2C12 cells cultured in PCL/rGNP@-10% scaffolds. H) Viability assay 24 hours after seeding and after 8 days in culture. All scale bars: 200 μm; green: myosin heavy chain (MHC); blue: DAPI.

Compared to 2D cultures (Figure 5C), cells grown on PCL, PCL/rGNP@-2%, and PCL/rGNP@-10% scaffolds embedded in Matrigel/collagen constructs show highly organized myotubes that align along the shape of the MEW scaffolds (Figures 5D,F,G and S6). All three scaffold compositions performed equally well in terms of differentiation (Figure 5E). However, PCL scaffolds performed significantly better for myoblast-to-myotube fusion and showed highly organized myotube alignment compared to the magnetized PCL scaffolds. It needs to be considered that all these experiments have been performed under static conditions. To evaluate the effects of dynamic cultures in response to a magnetic field, further experiments need to be performed. Furthermore, to assess scaffold biocompatibility, a viability assay was performed. A significant difference 24 h after seeding in cell viability was found between PCL, PCL/rGNP@-2%, and PCL/rGNP@-10%. On the contrary, no significant difference in cell viability after 24 h was found between PCL/rGNP@-2% and PCL/rGNP@-10%. The C2C12 cells were differentiated for 6 days amounting to a total culture time of 8 days. No significant difference in cell viability after 8 days of culture was found between PCL, PCL/rGNP@-2%, and PCL/rGNP@-10% scaffolds. Our results show that, although the PCL/rGNP@ scaffolds initially show a lower cell viability, long-term culture for a total of 8 days does not show a significant difference in cell viability compared to C2C12 cells in PCL scaffolds. Overall, both PCL/rGNP@-2% and PCL/rGNP@-10% scaffolds showed similar performance and can therefore be used for future studies. Moreover, C2C12 myoblast-laden MEW/Matrigel/collagen constructs with PCL/rGNP@-10% composition were able to successfully undergo reversible bending while immersed in culture medium under cyclical magnetic field loading (Video S1). Altogether, these observations suggest that these magneto-active constructs can allow for cyclical mechanical stimulation and guide cell orientation with high viability.

Importantly, to date, relatively few methods exist for recapitulating the fiber anatomy of skeletal muscle. Differentiation protocols of myoblasts such as C2C12 in 2D culture are relatively effective on gene expression level. However, morphologically, C2C12 cells differentiated in 2D do not represent physiological myotubes. Here, we demonstrated that, in a MEW fiber/hydrogel platform, C2C12 cells can be guided to organize themselves along scaffolds. In addition, compared to 2D, C2C12 cells were able to form myotubes that more closely resemble those *in vivo*. Furthermore, the positive effects of magnetic stimulation on myoblasts have been shown in multiple studies both *in vitro* and *in vivo*, as summarized recently by Mueller *et al*. ^[56]^. While most studies focus on uniaxial alignment, our approach is innovative as MEW provides flexibility to create complex architectures that allow for highly organized constructs in which alignment is multi-directional.

### 5. Future perspectives on integration of multi-material scaffolds towards complex macroscale geometries

The flexibility here described for fiber deposition *via* MEW was leveraged for the future development of scaffolds that tailor the mechanical and responsive properties of cellular constructs to mimic those of live tissues, especially skeletal muscle. Here, we showed the design potential of MEW PCL/rGNP@ scaffolds based on grid geometry and zonal distribution of active/nonactive materials (Figure 6). Firstly, the choice of grid geometry might allow to approximate diverse cell orientations, for instance, based on a cartesian grid for linear orientation profiles or a polar grid for circular orientations. These complex geometries could help emulating the wide range of cellular configurations found across diverse skeletal muscle types, e.g. parallel and circular muscle. Second, fibers composed of diverse materials can be differentially deposited stepwise *via* MEW, in order to generate scaffolds with active and nonactive zones that mimic tissue regions with different fiber stiffness and different magneto-responsive deformation. The adjacent deposition of PCL/rGNP@ and PCL yielded hybrid scaffolds consisting of magnetically active and inactive regions, which could be used as scaffolds with location-dependent deformation, for instance, as observed in muscle insertions. Overall, the design considerations here described for MEW of PCL and PCL/rGNP@ composites expand the current fabrication possibilities for mimicking the diverse structural configurations of skeletal muscle. Importantly, the newly developed magnetically deformable microfiber scaffolds will provide a platform for the further study of the role of magnetic fields on skeletal muscle *in vitro*. In particular, the use of patient induced pluripotent stem cell (iPSC)-derived skeletal muscle under dynamic stimulation triggered by magnetic fields remains relatively unexplored. Our platform opens new possibilities for disease modeling to better understand the effect of mechanical stimulation on muscle differentiation, as well as a disease modeling system for DMD and FSHD.

**Figure 6.**
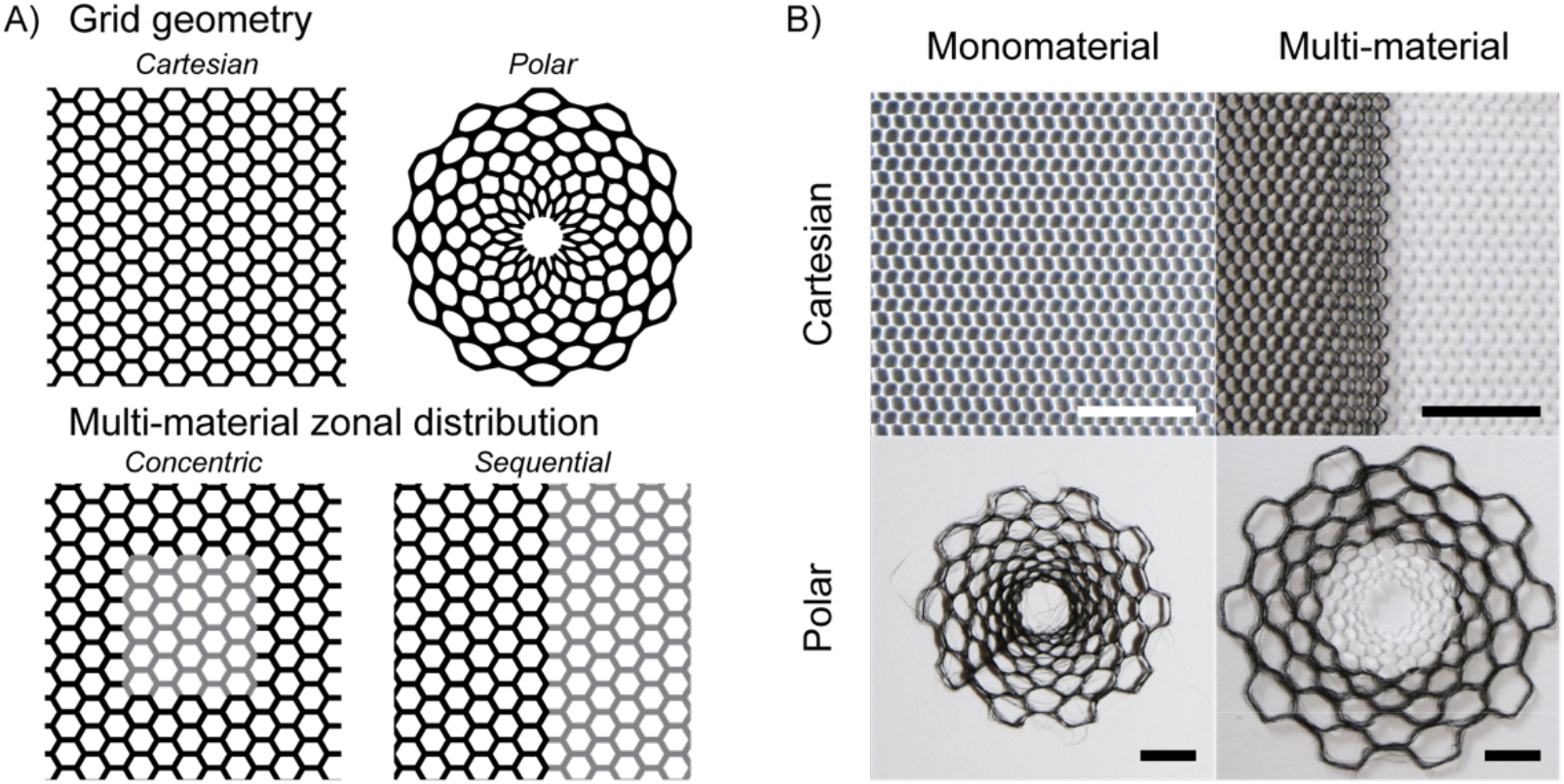
Integration of magneto-active MEW scaffolds based on multiple materials with complex fiber orientations. A) Design considerations for the integration of geometrically complex, differentially active MEW scaffolds (PCL/rGNP@ in black; PCL in grey). B) Scaffold grid geometry and zonal distribution of active material for introducing active/nonactive interfaces and complex magneto-triggered deformation profiles that recapitulate the diverse mechanical microenvironments (PCL/rGNP@-10% in black; PCL in white). Scale bars: 5 mm.

## Conclusion

In summary, we showedthe fabrication of magnetically deformable microfiber meshes with controlled hexagonal architectures via melt electrowriting of a cytocompatible material composite. When combined with Matrigel/collagen gels, these microstructured scaffolds were capable of undergoing reversible bending triggered by cyclical application of external magnetic fields. Moreover, these magneto-active constructs generated 3D culture environments that guided cell alignment along scaffold microfibers. We envision our novel approach as an innovative platform for the rational design of bio-inspired scaffolds that provide remote stimulation, thus advancing the *in vitro* modeling of disease and regeneration in skeletal muscle and other soft tissues.

## Acknowledgments

GC, JM, and MC acknowledge financial support from the Netherlands Organization for Scientific Research (NWO) through the Gravitation Program “Materials Driven Regeneration” (024.003.013) and the European Union Horizon 2020 program through project BRAV3 (874827). MC acknowledges the financial support from the NWO through project RePrint (OCENW.XS5.161). OD, FGS, and NG acknowledge the financial support from the Novo Nordisk Foundation (NNF21CC0073729), FSHDglobal, and the Stichting Utrecht Singelswim. JM, FDM, and AMP acknowledge financial support from the Portuguese Foundation for Science and Technology (FCT) / Ministry for Science, Technology, and Higher Education (MCTES PIDDAC) through projects ALiCE (LA/P/0045/2020) and LEPABE (UIDB/00511/2020 and UIDP/00511/2020) and the Institute for Research and Innovation in Health i3S (UIDB/04293/2020); through the UT Austin PT Program (project UTAP-EXPL/NPN/0044/2021); from FEDER funds through the COMPETE 2020–Operational Programme for Competitiveness and Internationalisation, Portugal; and from the Norte Portugal Regional Operational Programme (NORTE) within the Portugal 2020 Partnership Agreement of the European Regional Development Fund through project 2SMART (NORTE-01-0145-FEDER-000054). AMP acknowledges financial support from the FCT through the Scientific Employment Stimulus (Individual Call, CEECIND/03908/2017).

## Supplementary Materials

**Figure S1.**
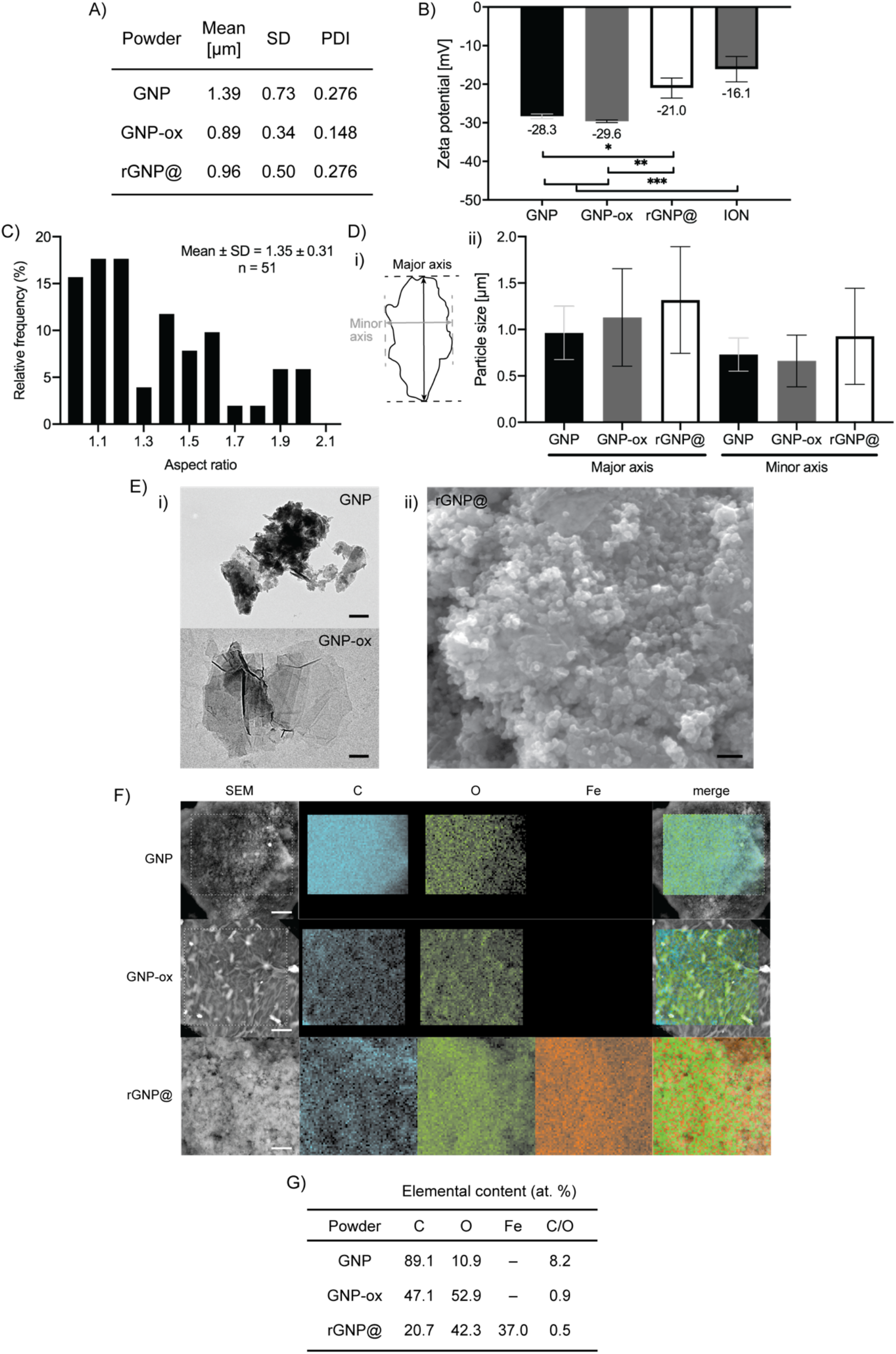
Morphology, particle size, and composition of graphene-derived and magnetized particles. A) Particle size distribution for GNP, GNP-ox, and rGNP@, as obtained by number-weighted DLS; mean, standard deviation (SD), and polydispersity index (PDI, calculated as mean^2^/SD^2^). B) Zeta potential of graphene-derived and ION particles (one-way ANOVA with Tukey multiple comparisons). C) ION particle aspect ratio as determined from TEM. D) i) Schematic of a particle showing its major (M) and minor (m) axes; ii) Particle size distribution as determined from TEM, showing mean ± SD. E) i) TEM and ii) SEM images of magnetized particles; scale bars: 200 nm. F) SEM-EDS images of magnetized particles showing elemental distribution; scale bars: 5 μm. G) Elemental composition of graphene-derived and magnetized particles obtained from EDS quantification.

**Figure S2.**
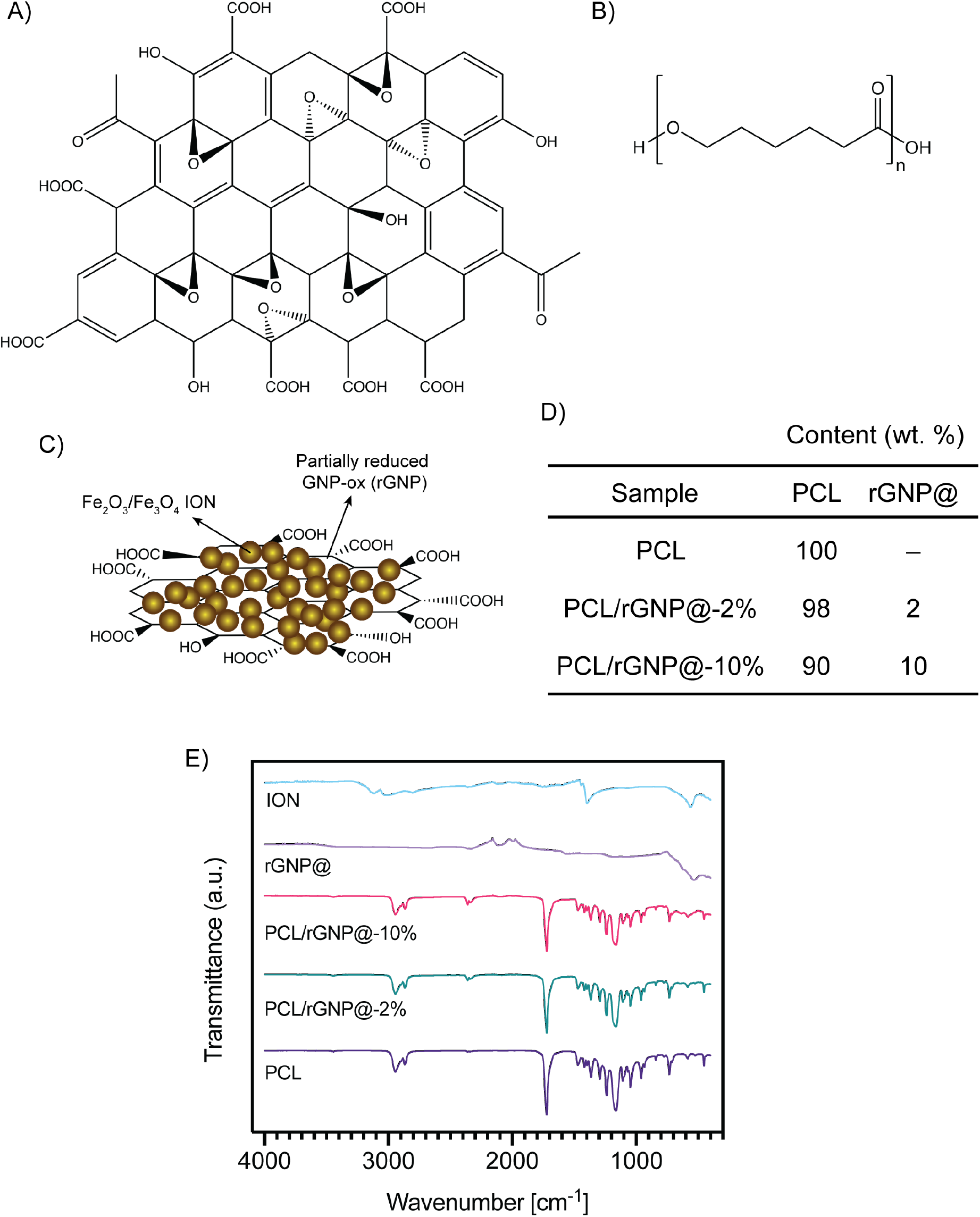
Composition of the PCL/rGNP@ composite. Chemical structures of A) oxidized graphene (GNP-ox) and B) poly-(ε-caprolactone) (PCL). C) Schematic structure of magnetic reduced graphene nanoplatelets (rGNP@), which consists of iron oxide nanoparticles (ION) obtained as a mixture of magnetite and maghemite deposited on partially reduced GNP-ox nanoplatelets. D) Nominal compositions of all PCL and PCL/rGNP@ composite samples. E) FTIR spectra of PCL, PCL/rGNP@ composites, and magnetic particles.

**Figure S3.**
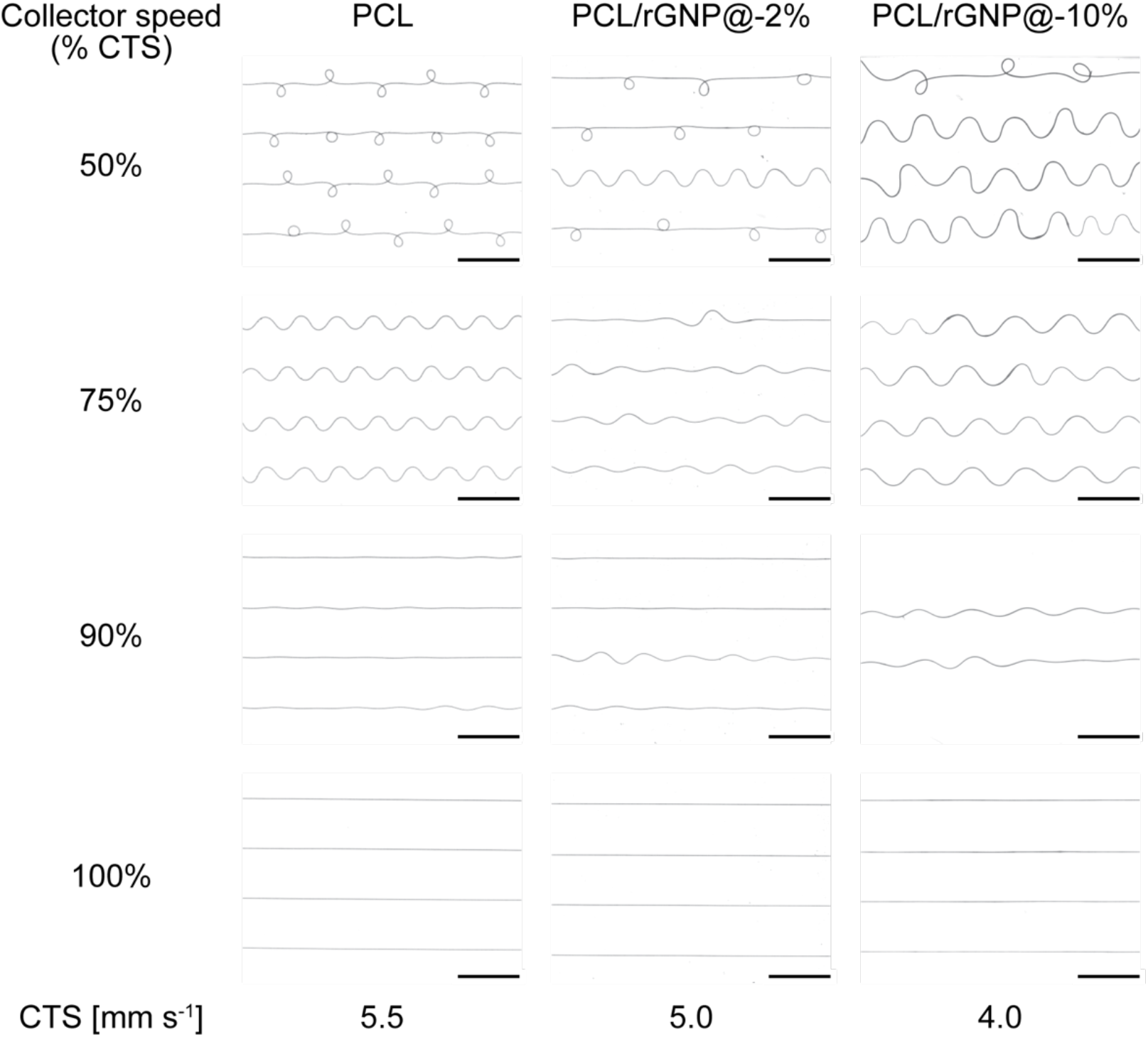
Effect of composition and collector speed on MEW fiber morphology. Scale bars: 1 mm.

**Figure S4.**
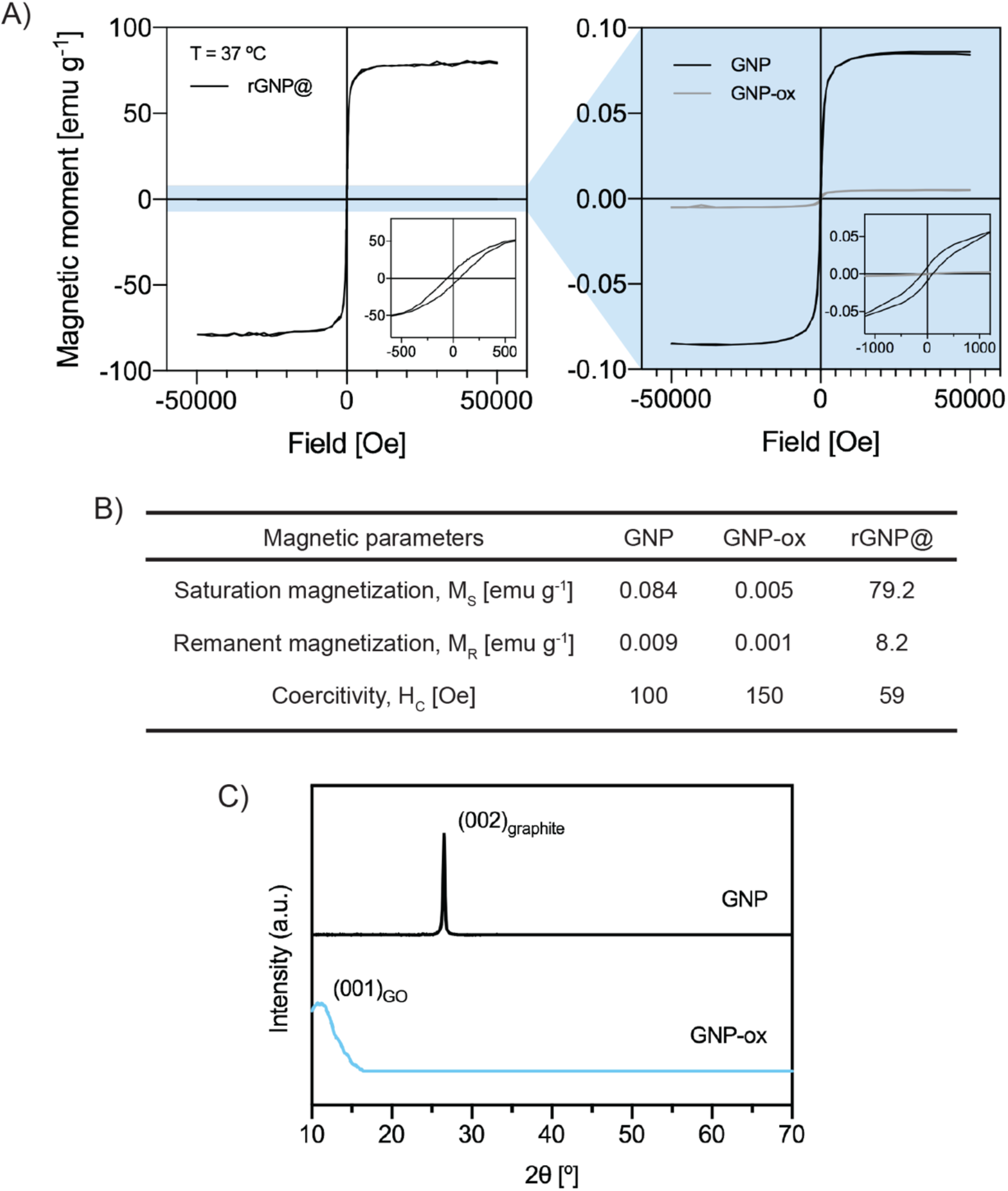
Magnetic and crystallographic properties of graphene-derived and magnetized rGNP@ powders. A) Magnetic hysteresis curves and B) magnetic parameters of magnetized powders at 37 °C. C) X-ray diffractograms of GNP and GNP-ox powders.

**Figure S5.**
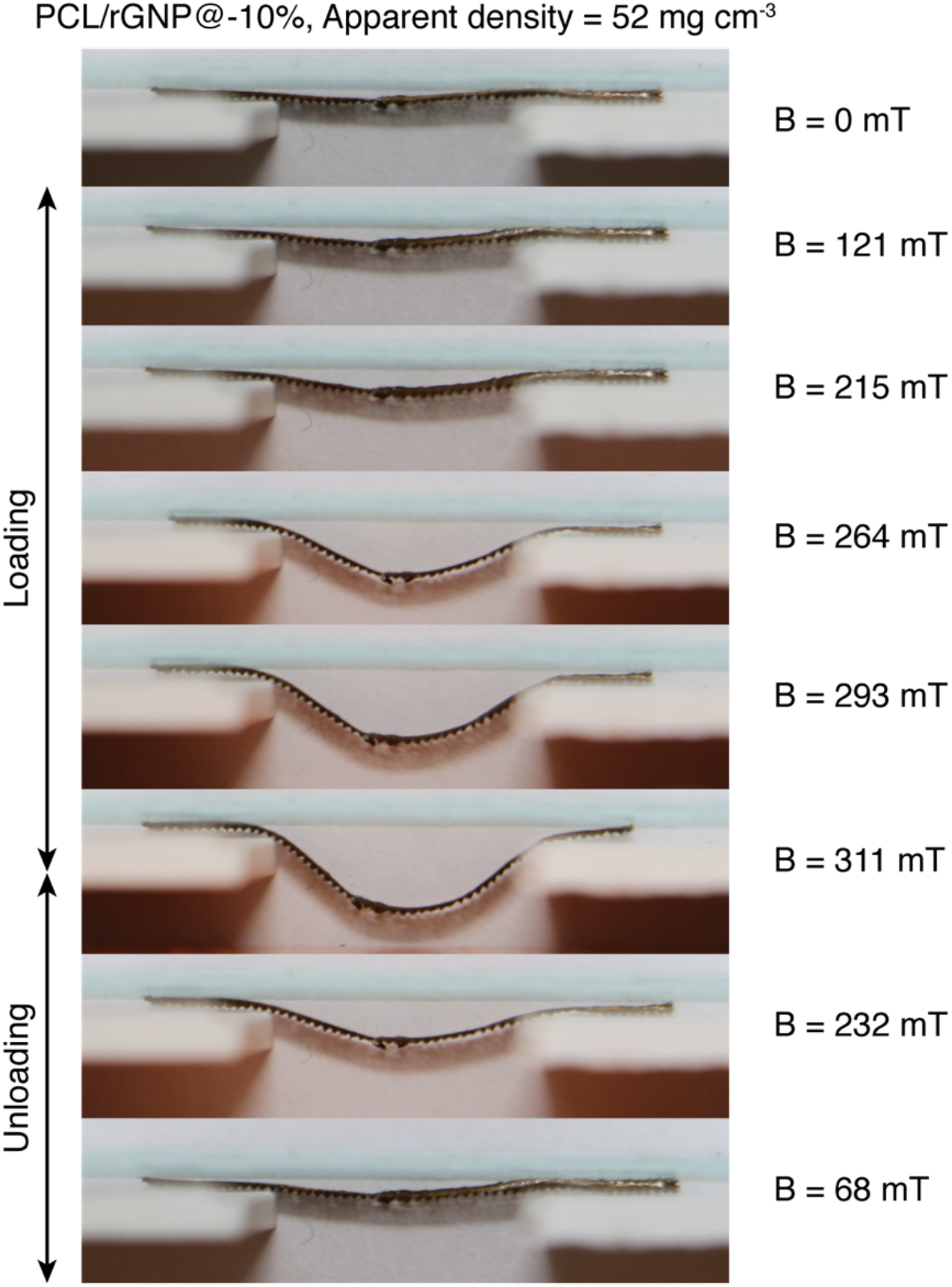
Snapshots of actuation test on a PCL/rGNP@-10% MEW scaffold under different applied magnetic flux intensities.

**Figure S6.**
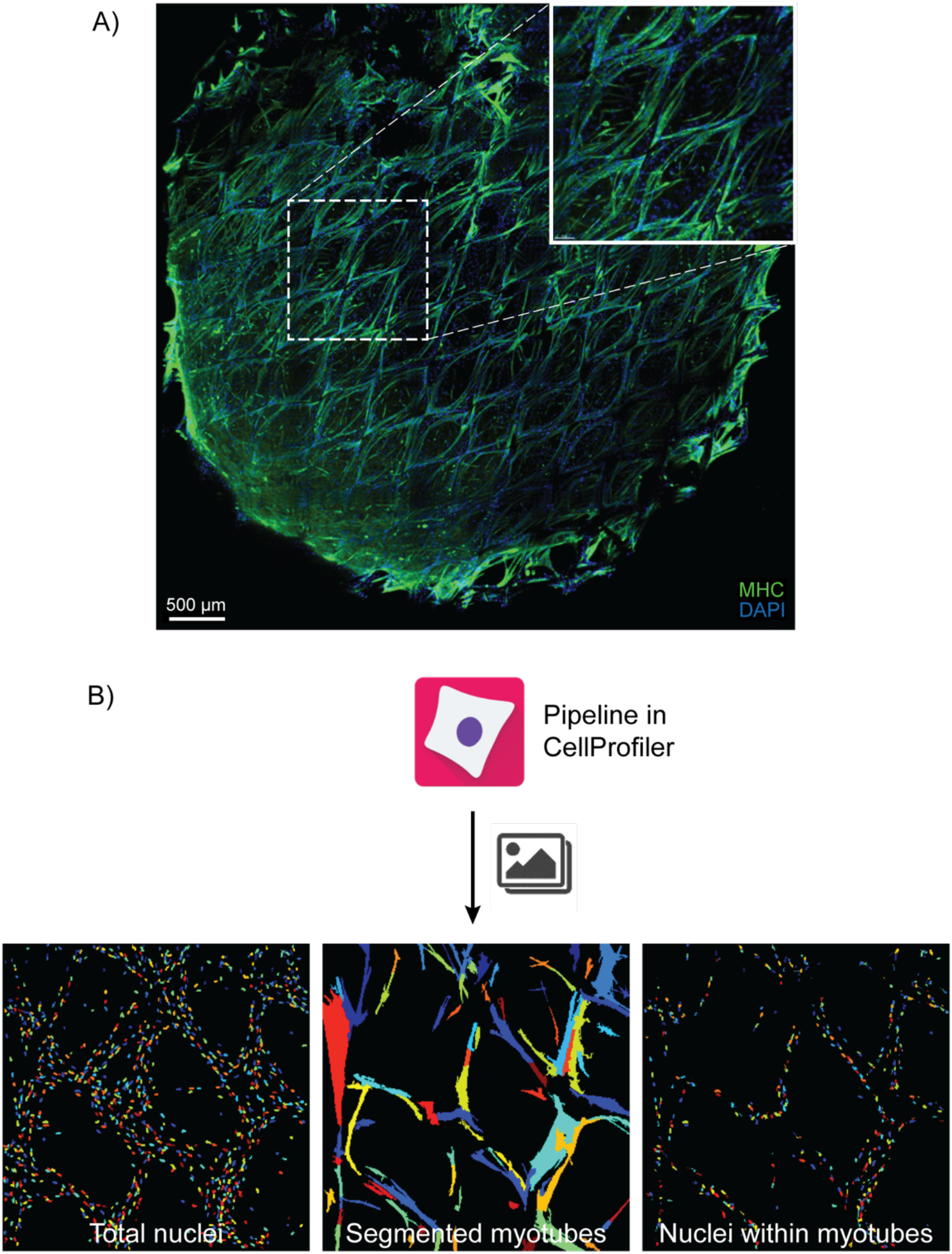
A) Confocal imaging of C2C12 cells cultured on whole MEW/Matrigel/collagen constructs; myosin heavy chain (MHC) = green; DAPI = blue. B) Workflow of image analysis for C2C12 cultures on whole constructs.

**Video S1**. Real-time cyclical bending of a C2C12 cell-laden MEW/Matrigel/collagen construct containing a PCL/rGNP@-10% scaffold, performed at 23 °C inside DMEM with an electromagnet that generated cycles of magnetic loading and unloading.

